# Selection of Fluorinated Aptamer Targeting RNA Element with Different Chirality

**DOI:** 10.1101/2024.06.24.600466

**Authors:** Yuliya Dantsu, Ying Zhang, Wen Zhang

## Abstract

The development of RNA aptamers with high specificity and affinity for target molecules is a critical advancement in the field of therapeutic and diagnostic applications. This study presents the selection of a 2’-fluoro modified mirror-image RNA aptamer through the *in vitro* SELEX process. Using a random RNA library, we performed iterative rounds of selection and amplification to enrich aptamers that bind specifically to the viral frameshift element which contains the opposite chirality. The unnatural chirality of the aptamer improved its enzymatic stability, and the incorporation of 2’-fluoro modifications was crucial in enhancing the binding affinity of the aptamers. After nine rounds of SELEX, the enriched RNA pool was sequenced and analyzed, revealing the dominant aptamer sequences. The selected 2’-fluoro modified mirror-image RNA aptamer demonstrated a dissociation constant of approximately 1.6 μM, indicating moderate binding affinity with the target and exceptional stability against nuclease degradation. Our findings highlight the potential of 2’-fluoro modified mirror-image RNA aptamers in enhancing the stability and utility of RNA-based therapeutics and diagnostics, paving the way for future applications in diverse biomedical fields.

---

RNA aptamers, short single-stranded RNA molecules capable of folding into versatile three-dimensional geometries and binding to specific target molecules with high affinity and selectivity, have emerged as versatile tools with numerous applications in biotechnology, medicine, and diagnostics^1, 2^. The selection of RNA aptamers with desired properties typically involves iterative rounds of *in vitro* evolution, a process known as SELEX (Systematic Evolution of Ligands by EXponential enrichment). SELEX allows for the identification of aptamers that can recognize a wide range of targets, including proteins, small molecules, and even whole cells, with high affinity and specificity^3^. Traditional RNA aptamers, however, are susceptible to degradation by nucleases, limiting their stability and utility in various biological applications^4, 5^. To date, tremendous efforts have been devoted targeting nucleic acid structures to engender the stable candidates in biological fluids^6^.

One effective strategy to overcome the limitation of stability in RNA therapeutics is to utilize mirror-image RNA aptamers, where the ribose sugars of the RNA backbone are replaced with L-ribose. This modification renders the aptamers resistant to nuclease degradation while preserving their binding properties. The mirror-image approach not only enhances the stability of RNA aptamers but also broadens the scope of RNA-based therapeutics and diagnostics. To date, L-nucleic acid-based aptamers have been successfully isolated to target a variety of disease-related elements, including small molecules^*7-10*^, non-coding RNAs^*11-16*^, amino acids^*17-21*^, and proteins^*22-27*^. These L-aptamers have demonstrated exceptional affinities and significantly improved stability, binding selectively to their targets through unique tertiary structural features and effectively blocking disease-related biological processes. Given their remarkable stability and high affinity, L-aptamers represent a promising avenue for the development of advanced therapies. Their unique properties could be harnessed to create next-generation RNA-based treatments that are not only more stable in biological environments but also highly specific to their molecular targets. As such, L-aptamers hold significant potential for addressing a wide range of diseases, making them an exciting focus for future therapeutic development.

In this study, we report the selection and characterization of a new mirror-image RNA aptamer using *in vitro* SELEX. The target molecule for aptamer selection is the attenuator hairpin structure of coronaviruses, a critical regulatory element that controls the frameshift mechanism of virus gene expression. All coronaviruses, including severe acute respiratory syndrome (SARS)-CoV-2, utilize the essential programmed -1 ribosomal frameshifting (−1 RF) strategy to regulate the RNA genome expression in the 2 overlapping translational open reading frames (ORF 1A and ORF 1B, Figure 1A)^28, 29^. In the -1 RF mechanism, the cis-acting elements in the mRNA can direct elongating ribosomes to shift the reading frame by 1 nucleobase toward the 5′-direction^30^. The stable and structured RNA frameshifting element (FSE) controls the unique -1 RF. FSE is composed of three elements, including attenuator hairpin (AH), slippery site and pseudoknot (Figure 1A). They can collectively function to prohibit the initiation of frameshifting, thereby altering the protein coding on viral mRNA. Among FSE RNA structures, the attenuator hairpin is located immediately 5′ of the slippery site, and it regulates -1 RF by attenuating its activity^31^. The primary sequence and predicted secondary structure containing AGCU tetraloop and bulges are shown in Figure 1B^32, 33^.

**Figure 1.**
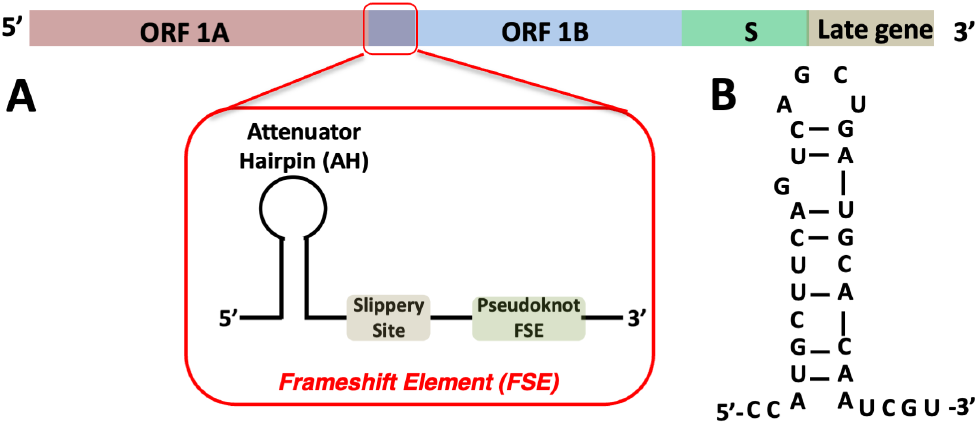
(A) Schematic representation of SARS-CoV-2 RNA genome. The location of the viral frameshift element was indicated by the red box. (B) The sequence and predicted secondary structure of the attenuator hairpin, which is the SELEX target.

Therefore, the controlling of gene expression by FSE is critical for viral replication and pathogenesis^34^. An effective strategy to disrupt normal viral gene expression involves using complementary strands, such as antisense oligonucleotides, to block the activity of the frameshift element (FSE) through RNase H-mediated degradation. However, antisense oligonucleotides must be in the native D-form to hybridize with viral RNA, which introduces the risk of undesired degradation of the therapeutic agent. Instead, another approach to inhibit the viral frameshifting and interfere the viral replication is to develop the structure-specific binder to target the FES element. There have been drug-like small molecules identified to selectively bind to the AH structure and impair the frameshifting of virus^32, 35^. These studies demonstrated the promise that the RNA genome of SARS-CoV-2 could be an ideal drug targets to disrupt the viral cellular functions. The future development of the structure-specific ligands to target virus genome could be based on small molecules or other macromolecules, which has the potential to serve as the complementary strategy of COVID-19 vaccine. Overall, by targeting this regulatory element with RNA aptamers, it is expected to modulate frameshifting dynamics and disrupt the production of essential viral proteins, thereby inhibiting viral replication and reducing viral pathogenicity.

In addition, fluorination is a prevalent chemical derivatization extensively utilized in pharmaceutical chemistry. This is attributed to the distinctive biophysical and biochemical properties of fluorine, such as its small atomic size and high electro-negativity^36^. Particularly, when replacing the H or OH group by fluorine element in nucleic acid drugs, the small size of fluorine leads to minimal steric effect on nucleic acid functions, and the significant electronic effect of fluoride modification could lead to the characteristic pseudo hydrogen bonding, as well as the alteration of nucleic acid sugar pucker and overall structures^37^. Thus, fluorination has become popular approach in nucleic acid-based therapeutics, including anticancer and antiviral nucleosides^38^, and fluoride-oligonucleotides as therapeutic antisense^39^, and siRNA^40^ drugs. In particular, 2′-F-modification has been introduced to therapeutic aptamer selections, in which fluorination contributed to the enhanced binding and biological stability of the aptamers^41-43^. In our lab, the 2′-F-derivatized L-pyrimidines have been successfully synthesized^44, 45^, and it was discovered that the fluoride atom contributed to rigidify the L-ribose to enhance the helical structural and thermal stability^46^, engender the pseudo-hydrogen bonds and strengthened stacking interactions^47, 48^. Therefore, we attempt in this study to further expand the application of 2′-fluoro-L-nucleic acids by isolating the functional mirror-image aptamer, which has enhanced stability and binding affinity, thus improving its potential as a therapeutic agent against viral infections.

Due to the lack of DNA and RNA polymerases capable of recognizing mirror-image molecules, our SELEX was completed in a mirror-image manner (Figure 2A). In order to select L-aptamers that can bind to naturally occurring D-type RNA, the target molecule in the actual SELEX was L-type, while the aptamers selected in each round were based on D-type. Therefore, the biotin-labeled L-type target molecule, L-RNA attenuator hairpin (L-AH, Figure S1), was synthesized by solid-phase synthesis and used as the target molecule in each round, tightly binding to streptavidin molecules on magnetic beads to select functional aptamer sequences. Random DNA library was synthesized by IDT, and the random region was composed of 45 nucleotides flanked by 5’ and 3’ primer regions (Figure 2B). At the primer regions, T7 promoter binding sequence was included for the *in vitro* transcription. The 5′ and 3′ primer for PCR and *in vitro* transcription was 5′-TTCTAATACGACTCACTATAGGTTACCAGCCTTCACTGC-3′ and 5′-GTGTGACCGAC CGTGGTGC-3′, respectively. In the first round, 1 nmol random RNA library was prepared by *in vitro* transcription, using the Y639F T7 RNA polymerase mutant (Figure S2). In each round, the streptavidin coated dynabeads were first incubated with 0.1 mg/ml yeast tRNA for 2 hours at room temperature to block the nonspecific binding. The reaction mixture containing the RNA library and the corresponding buffer conditions was heated at 80 °C for 2 min, then slowly cooled to room temperature to form the RNA structure. The negative selection was first performed by incubating the RNA library with the prepared dynabeads for certain time (Table S1, 5 min in round 1, and increase in each round after). The supernatant was extracted and used for the following positive selection. In the positive selection, the 5′-bio-tinylated L-AH was added to the reaction mixture and incubated with the RNA library for certain time (Table S1, 60 min in round 1, and decrease in each round after). Then, the reaction was added to a 1 mg of the prepared dynabeads and incubated for additional an hour. The supernatant was discarded, and the dynabeads were washed twice using the washing buffer. The captured RNAs were eluted with 100 µl of elution buffer containing 25 mM NaOH and 1 mM EDTA, followed by neutralization with Tris-HCl buffer (pH 7.5) and purification with NEB RNA clean and concentration column. The process of selection and amplification was then repeated with the selection stringency progressively increased each round by employing longer wash times and decreasing template concentrations and incubation times (Table S1).

**Figure 2.**
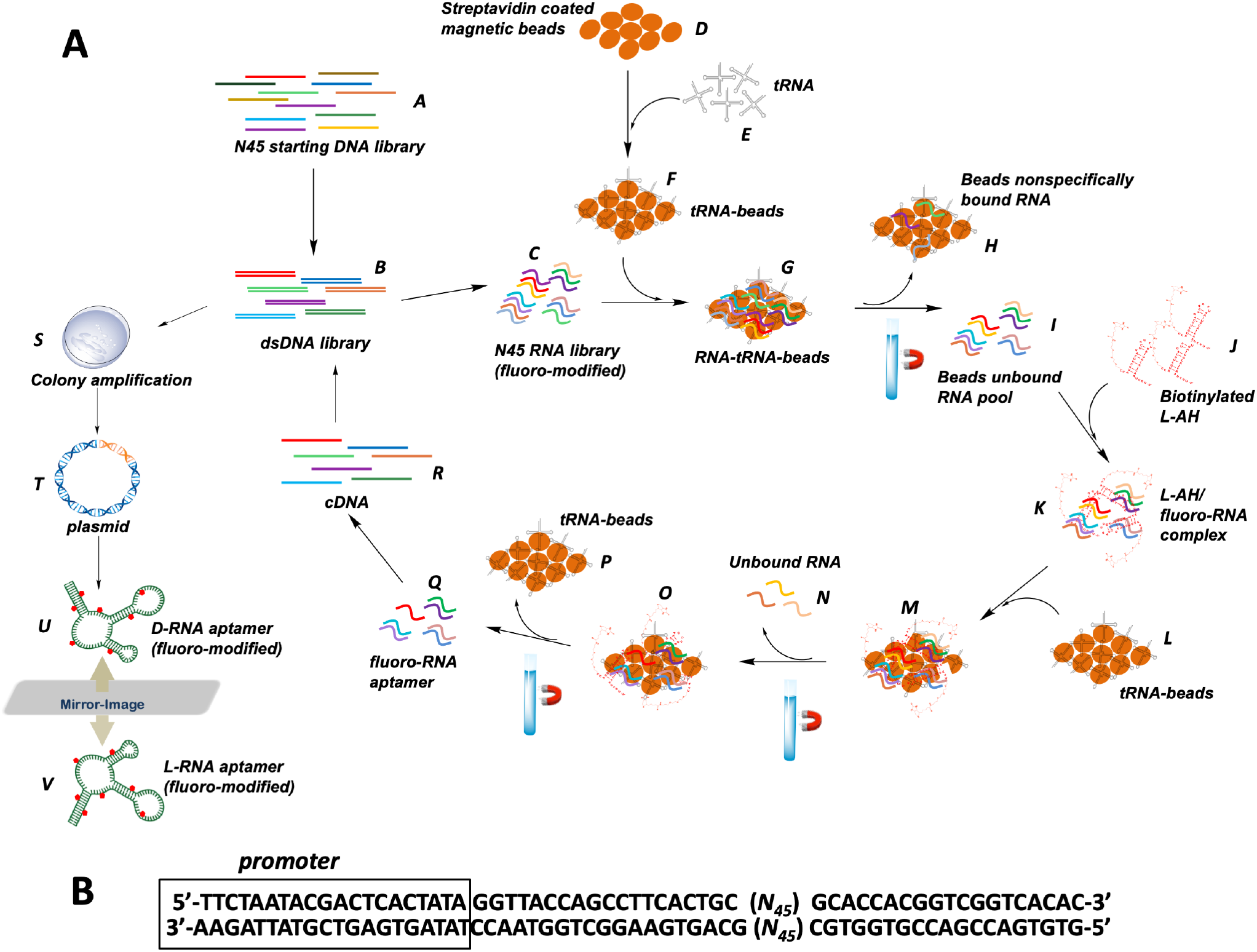
(A) Schematic representation of the mirror-image SELEX. (B) The sequence of dsDNA library in each round. The promoter region for T7 RNA polymerase recognition was boxed.

The enriched RNA library was then reverse transcribed using NEB ProtoScript^®^ II Reverse Transcriptase. After reverse transcription, the RNA library was degraded in a 50 µl solution which contained 0.2 M NaOH solution and was incubated at 95 °C for 10 min. The obtained cDNA single-stranded template was PCR amplified using the forward and reverse selection primers, and NEB Q5^®^ High-Fidelity DNA Polymerases. The subsequent dsDNA template was used for the mutant T7 *in vitro* transcription for the next round. Overall, we did nine rounds of selection in this study, and analyzed the SELEX result by High-throughput Sequencing in Purdue Genomics Core Facility.

From the sequencing results, six highly enriched representative sequences were identified (Figure 3A). The sequences and percentages of these candidates are shown in Table S2. Microscale thermophoresis (MST) binding assays were conducted to measure the binding affinity of the aptamers to the synthesized 5′-FAM labeled L-AH, with the K_d_ of Apt. 9.1 determined to be 1.4 µM (Figure S3). We also assessed the binding affinities of the other five most abundant sequences, identifying two additional aptamers with similar binding affinities: Apt. 9.2, with a Kd of ∼2.5 µM, and Apt. 9.3, with a Kd of ∼2.1 µM (Figure S4 and S5). The other three sequences, Apt. 9.4, 9.5, and 9.6, did not exhibit detectable binding affinities, despite their abundance in the sequencing results (Table S2). Using the RNA secondary structure prediction program Mfold^49^, the secondary structures of all six sequences were predicted. Aptamers 9.1, 9.2, and 9.3 were predicted to have possible cloverleaf-like structures, consisting of 83 nucleotides, 4 stems, and 3 loops (Figure 3B). The 2′-fluoro-modified uridine and cytidine residues are present in all stems and loops. In contrast, Mfold did not predict cloverleaf-like structures for Apt. 9.4, 9.5, and 9.6, which instead formed significantly different secondary structures, as shown in Figure 3B. It is also noted that the stem size in Apt. 9.1, 9.2, and 9.3 is all different, especially the lengths of stem 3 and stem 4. Considering that these three aptamers show similar modest binding affinity, it is suggested that the folded conformation of the aptamers, particularly the formation of three loops, is more crucial for its binding ability than the lengths of the stems in the structure. The cloverleaf-like structure observed in Apt. 9.1, 9.2, and 9.3 may play a key role in facilitating the high binding affinities of these sequences. In contrast, the lack of such a structure in Apt. 9.4, 9.5, and 9.6 likely contributes to their undetectable binding. This highlights the importance of secondary structure in aptamer functionality and provides valuable insights for future aptamer design, where structural features could be optimized to enhance binding affinity and specificity.

**Figure 3.**
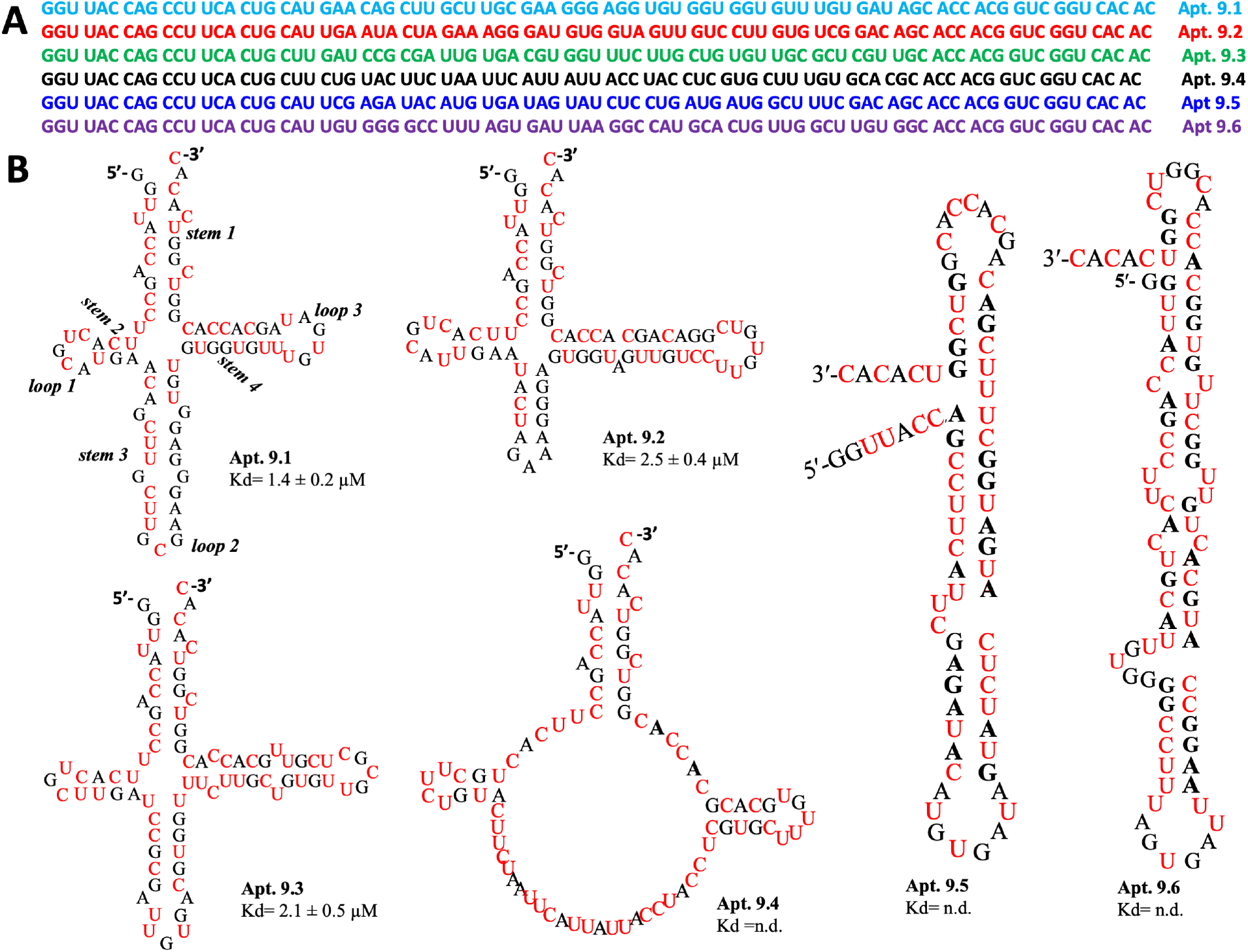
(A) The sequences of the most enriched aptamers after nine rounds of *in vitro* selection. (B) The predicted secondary structures of the most abundant sequences Apt. 9.1, 9.2, 9.3, 9.4, 9.5 and 9.6. In all the secondary structures, fluoro modified resides are labeled by red color, and the K_d_ values are listed below the structures.

We then focused on Apt 9.1 (Figure 4A) to do in-depth analysis about the aptamer’s binding features. The 5′ and 3′ primer regions of the aptamer were initially designed as fixed sequences for PCR amplification and T7 transcription purposes. However, when we attempted to truncate the primer regions to create a shorter aptamer, the binding affinity was completely lost (Apt 9.1-x, Figure S6). Secondary structure predictions indicated that truncating the primer regions caused the RNA to preferentially fold into a short hairpin geometry with 5′ and 3′ overhangs (Figure 4B). This distinct structural alteration likely accounts for the observed loss of binding affinity. Despite this challenge, we successfully obtained a 71-nucleotide truncated variant, Apt. 9.1-b, by selectively deleting bulges and flanking nucleotides, which retained the original K_d_ of 1.4 µM (Figure 4C and S7). To investigate which loop/stem regions contributed to the aptamer binding, we then synthesized 3 truncated aptamers (Figure 4D, Apt. 9.1-c, 9.1-d and 9.1-e, Figure S8), and one loop/stem section was deleted in each aptamer mutant. It was discovered that, when loop 1 was truncated, Apt. 9.1-c presented much weaker binding affinity (K_d_ >100 µM). When loop 2 or loop 3 was truncated, no binding affinity could be measured in Apt 9.1-d and 9.1-e. This result indicates the importance of all three RNA loops, which may fold into compacted three-dimensional conformation by interacting each other. To further validate our MST measurements of binding affinity, we conducted a gel shift assay by titrating the FAM-labeled L-AH with D-Apt. 9.1-b. The results of the electrophoresis are presented in Figure S9. The K_d_ value determined from this assay was approximately 1.9 µM, which is slightly higher than the K_d_ value of 1.4 µM obtained from the MST measurement. This discrepancy in binding affinities is likely due to the inherent differences between the two methods. The MST assay measures binding affinity in solution, which may more accurately reflect the true binding conditions. In contrast, during the prolonged gel electrophoresis process, the RNA may become partially denatured, potentially affecting the binding interaction. Therefore, the K_d_ value of 1.4 µM determined by MST is likely a more accurate representation of the true binding affinity.

**Figure 4.**
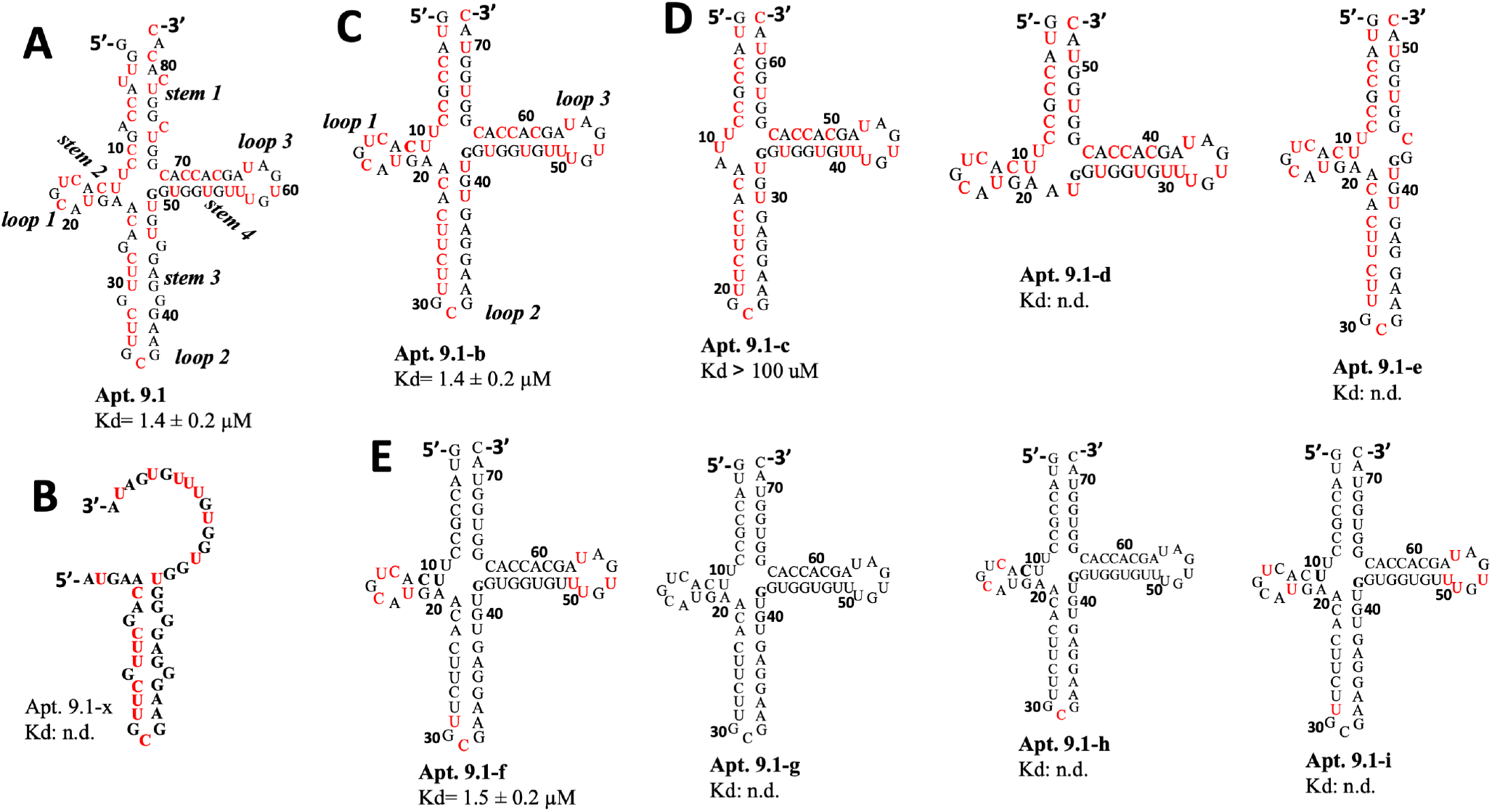
(A) The predicted secondary structure of Apt. 9.1. (B) The predicted secondary structure of Apt. 9.1-x, which has the primer regions truncated. (C) The predicted secondary structures of Apt. 9.1-b. (D) The predicted secondary structures of truncated versions of Apt. 9.1-b. (E) The predicted secondary structures of the functional aptamer Apt. 9.1-f, 9.1-g, 9.1-h, and 9.1-i, which contains limited number of fluoro modifications. In all the secondary structures, fluoro modified resides are labeled by red color, and the Kd values are listed below the structures.

One more important question is whether the fluoride modifications contribute to the aptamer folding and binding, due to its unique electronegativity and ability of forming inter-and intramolecularly hydrogen bonds. We then chemically synthesized the mutated chimeric aptamer Apt. 9.1-f, which contained the 2′-fluoride modified cytidine and uridine residues in all three loops, and the wild-type uridine and cytidine residues in the stem regions (Figure 4E). The K_d_ of this aptamer was measured to be 1.5 µM (Figure 5A, black), which suggests that the 2′-fluoro-modifications in the stem regions don’t present significant contribution to the RNA folding and aptamer binding. The function of those fluoride atoms were likely the modest enhancement of thermostability of RNA duplex, which had been observed before^44, 45^. We then attempted to replace all the 2′-fluoro-pyrimidines with native residues, and found the aptamer mutant, Apt. 9.1-g, entirely lost the binding affinity. Similarly, when only 2′-fluoro modified cytidines or 2′-fluoro modified uridines were introduced in the aptamer (Apt. 9.1-h and Apt. 9.1-i), no binding affinity could be detected, which indicated the importance of those 2′-fluoro modifications. When the wild-type 2′-OH groups were replaced with 2′-F atoms, the original hydrogen bonding patterns likely changed and the electronegativity of fluoride could cause local disturbance of sugar pucker conformation, all leading to the destabilization of the aptamer cloverleaf structure.

**Figure 5.**
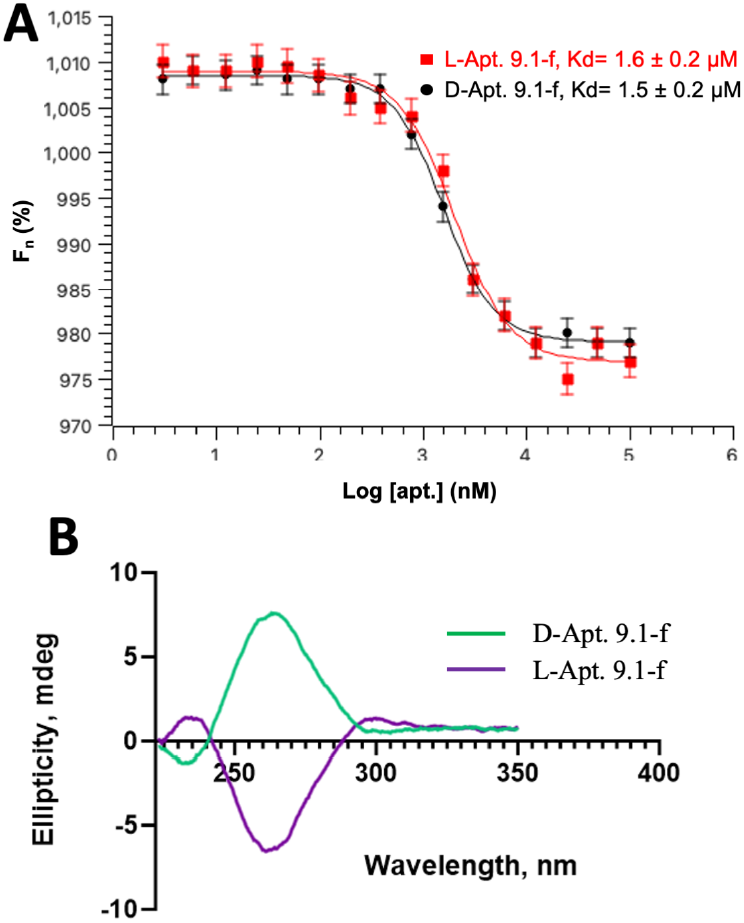
(A) Binding curve in MST assays generated from the difference in initial fluorescence intensity compared to the intensity in the presence of heat. D-Apt. 9.1-f against FAM-L-AH (black), and L-Apt. 9.1-f against FAM-D-AH (red) are presented. The error bars represent the standard deviation of 3 independent replicates. (B) Normalized CD spectra of D-and L-type aptamers Apt 9.1-f.

In our SELEX experiment, 10 mM MgCl_2_ was used to facilitate aptamer folding and its binding to the AH element. However, given the low concentration of Mg^2+^ under physiological conditions, it is important to investigate whether the aptamer can effectively bind to AH at lower Mg^2+^ concentrations. We measured the binding affinity of Apt 9.1-b under conditions of 2 mM and 0 mM Mg^2+^. With 2 mM Mg^2+^ present, the K_d_ of the aptamer increased dramatically to over 100 µM, and in the absence of Mg^2+^, the binding affinity was entirely lost (Figure S10 and S11). These results suggest that Mg^2+^ is crucial for maintaining the aptamer’s structure, enabling it to bind to its target. These findings emphasize the critical role of Mg^2+^ in stabilizing the aptamer’s structure for effective binding, which is a key consideration for its potential therapeutic application. The significant drop in binding affinity under physiological Mg^2+^ conditions indicates that further optimization is necessary to develop an aptamer that retains its binding capability at lower Mg^2+^ concentrations. Addressing this challenge could enhance the practical applicability of the aptamer in therapeutic settings.

We then investigated and compared the binding affinities between D-Apt. 9.1-f with L-AH versus L-Apt. 9.1-f with D-AH. We found them to be similar to each other, with the K_d_ to be 1.5 ± 0.2 and 1.6 ± 0.2 µM (Figure 5A). The difference between the two binding affinities could be due to the different qualities of chemicals in the solid-phase syntheses. In contrast, when the aptamer was used to treat the AH target with the same chirality (L-aptamer targeting L-AH, or D-aptamer targeting D-AH), no binding affinity could be detected. The results indicate the feature of cross-chiral recognition of our selected aptamer, as well as its specificity when binding to target.

We then performed the circular dichroism studies to characterize the chiralities of the two aptamers. The corresponding spectra are shown in Figure 5B. The data revealed that the D-form 2’-F-modified Apt. 9.1-f adopted a structure containing a high positive band at 266 nm and a weak negative band at 236 nm, which was generated by the characteristic A-form RNA duplexes in the aptamer^50^. In contrast, the chiral inversion of L-Apt. 9.1-f displayed a high negative band at 266 nm and a weak positive band at 236 nm. The data illustrates the two synthetic aptamers have the mirror-image conformations as the chiral counterparts.

One significant advantage of using L-nucleic acids as therapeutic agents is their biological stability when circulating in vivo. To assess the stability of the L-aptamer, we exposed L-apt. 9.1-f to 25% human serum and found that the L-aptamer remained intact for 24 hours. In contrast, the D-aptamer, despite having fluoride modifications on the RNA backbone, was completely degraded within 16 hours (Figure 6A). Additionally, we tested whether the aptamer could be efficiently delivered into cells using a traditional transfection reagent. As a proof of concept, we selected MDA-MB-468^51^ cells as a model and successfully delivered Cy5-labeled L-apt. 9.1-f into the cell culture using Lipofectamine 3000 reagent. The internalization of L-apt. 9.1-f was confirmed by the Cy5 signal observed inside the cells, with the signal localized around the nuclei, indicating that the aptamer molecules were delivered into the cytoplasm rather than the nuclei (Figure 6B). Furthermore, treatment with the L-RNA aptamer did not result in any reduction in cell growth, demonstrating the in vitro biosafety of using L-RNA as a therapeutic agent. To assess the potential immune response that L-RNA might trigger, we conducted an immune response analysis by measuring the expression level of nuclear factor kappa B (NF-κB). It is well-established that double-stranded RNA can activate Toll-like receptor 3 (TLR3), which in turn stimulates NF-κB family members to regulate the transcription of cytokines and antimicrobial effectors, playing a crucial role in both innate and adaptive immune responses^52, 53^. When HEK293t cells were treated with 5 µM L-apt 9.1-f and D-apt 9.1-f, no significant immune response was observed compared to the vehicle group (Figure 6C). This result indicates that the use of L-RNA aptamers as molecular therapies is likely safe. We did not observe a significant difference between L-and D-RNA, possibly due to the small size of these aptamer molecules, which may be insufficient to activate TLR3. These findings underscore the potential of L-aptamers as robust and stable therapeutic candidates, capable of resisting degradation in biological environments and being effectively delivered to target cells. It also provides preliminary evidence supporting the biosafety of L-RNA aptamers, which is a critical consideration for the development of RNA-based therapies, as immune activation can lead to adverse effects. This Moving forward, we plan to conduct a more comprehensive investigation into the immune response triggered by L-RNA therapies, further ensuring their safety and efficacy in clinical applications. This research lays the groundwork for the safe use of L-RNA in therapeutic settings, offering a promising path forward for the development of innovative molecular treatments.

**Figure 6.**
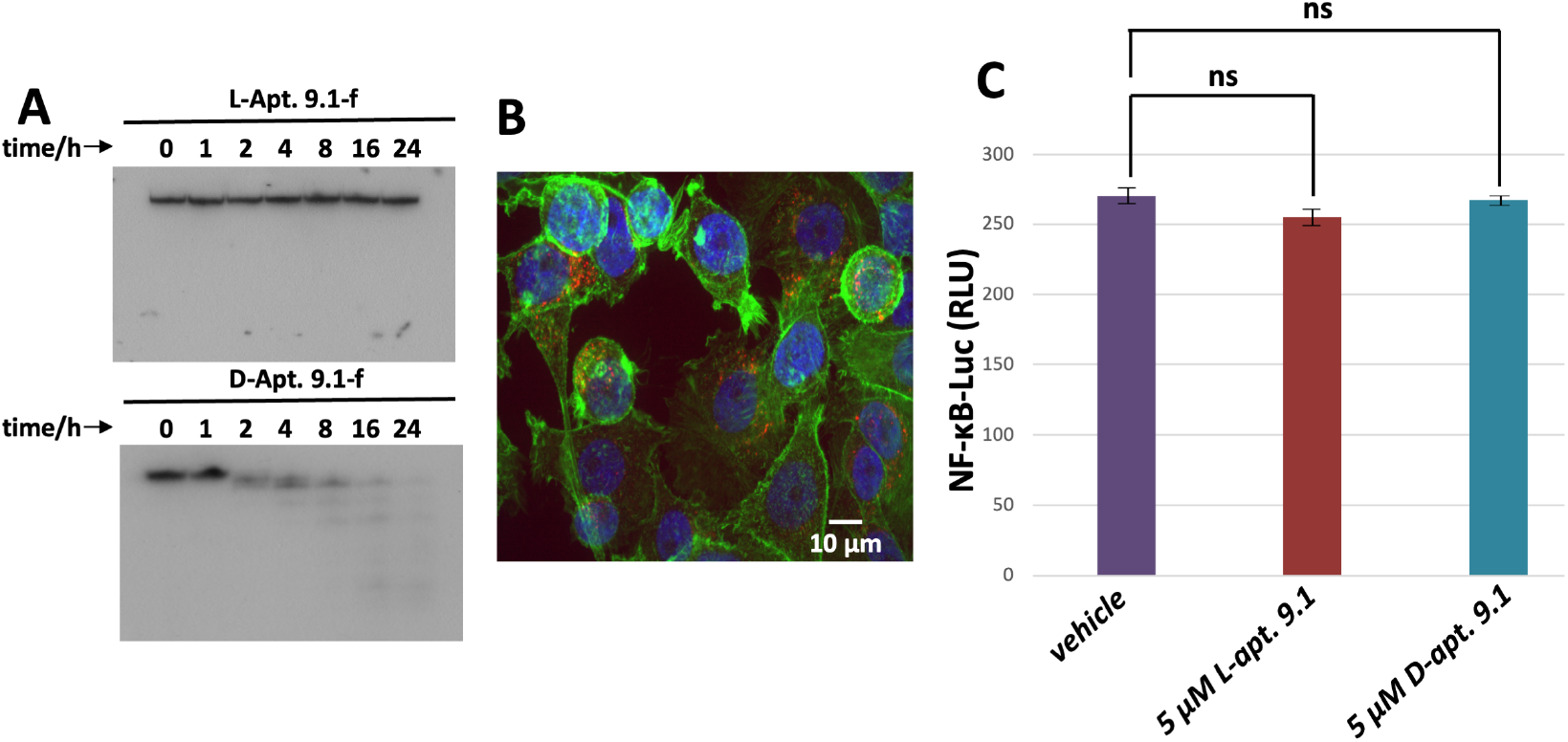
(A) Serum stability assay of L-and D-aptamers over 24h. (B) Confocal microscope assay showing the delivery of L-aptamer to MDA-MB-468 cells (blue: nuclei; green: cytoskeleton; red: Cy5 labeled L-aptamer). (C) Expression of NF-κB-regulated genes by L-and D-aptamers.

In this study, we successfully identified an L-RNA aptamer capable of binding to a native RNA element, demonstrating significant potential for therapeutic and diagnostic applications. Our findings indicate that the 2’-fluoro modifications within the L-RNA aptamer play a crucial role in its binding affinity, although the binding observed was modest rather than exceptionally strong. Moreover, comparative analysis with non-modified aptamers revealed that the introduction of 2’-fluoro groups improves binding strength and specificity towards the target RNA element, albeit not to the desired extent. These results align with previous studies that have shown the stabilizing effects of 2’-fluoro modifications in RNA-based therapeutics.

Given the modest binding affinity of our selected aptamer, further optimization of the SELEX conditions or the aptamer’s structure will be necessary for future therapeutic applications. One potential strategy is to modify the SELEX library by introducing additional constraints to enhance molecular stability and function. In the future, we may consider increasing the number of stem-loop regions in the library or designing a library rich in guanosine residues, which have been reported to improve binding diversity and affinity^54, 55^. Furthermore, we plan to investigate other modifications to the aptamer backbone, such as incorporating 2’-methoxy groups, which our lab has successfully synthesized in the past^56, 57^. Preliminary evidence suggests that 2’-methoxy modifications could offer greater stability and binding efficiency due to their enhanced resistance to nucleases and improved structural rigidity. Moreover, it is possible that some other sequences without fluoride modifications could offer even stronger binding affinity. Therefore, it is needed to further isolate wild-type L-aptamer and carefully compare its sequence, structure and binding affinity to fluoride-L-aptamer for the optimal drug development.

Additionally, it is important to note that while the *in vitro* binding characteristics are promising, *in vivo* efficacy and potential immunogenic responses need to be thoroughly evaluated in future studies. Additionally, exploring the structural dynamics of the L-RNA aptamer in complex with the native RNA element through advanced techniques such as X-ray crystallography or NMR spectroscopy could provide deeper insights into the binding mechanism.

Overall, our research highlights the potential of 2’-fluoro modified L-RNA aptamers as a foundational step towards developing robust tools for targeting native RNA elements. Further modifications, including the incorporation of 2’-methoxy groups, are anticipated to enhance these properties, paving the way for the development of novel RNA-based therapeutics and diagnostic agents.

## ASSOCIATED CONTENT

The Supporting Information is available free of charge at xxx

## AUTHOR INFORMATION

Notes

The authors declare no competing financial interest.

## ACKNOWLEDGMENT

We thank the Zhang lab for helpful discussions, insightful commentary and careful revision of the manuscript. We thank Dr. L. Zeng and the Chemical Genomics Core at IUSM for the help.

---

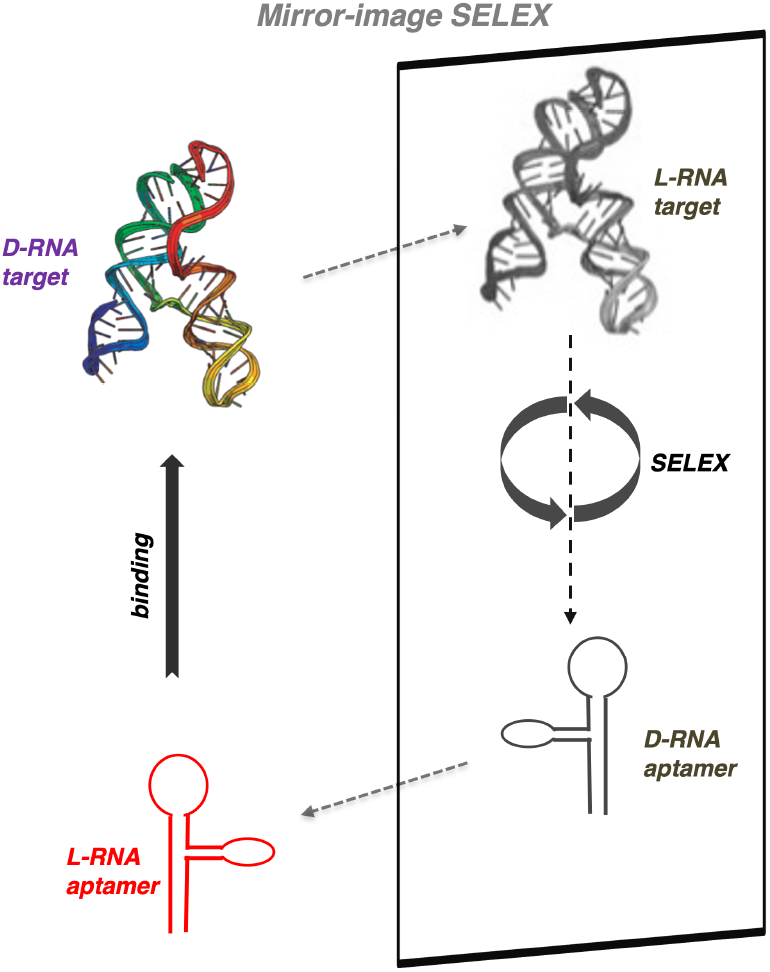
Insert Table of Contents artwork here

