## supporting information for "Selection of Fluorinated Aptamer Targeting RNA Element with Different Chirality"

Yuliya Dantsu, Ying Zhang and Wen Zhang*

*Department of Biochemistry and Molecular Biology, Indiana University School of Medicine, Indianapolis, IN 46202;
Melvin and Bren Simon Cancer Center, Indianapolis, IN 46202, USA

CONTENTS

1. General Methods ……………………………………………………………… 2
2. Protocols in SELEX, Enzymatic Reactions and Characterization of Aptamers… 3
3. References ……………………………………………………………………… 12

### 1. General Methods.

**1a. General considerations.** All chemicals were purchased from Fisher Scientific, VWR or Sigma-Aldrich unless otherwise noted. All the nucleoside phosphoramidites for solid-phase synthesis were from Chemgenes Corporation or Glen Research. All oligonucleotides, including primers used in selection cycles, aptamer candidate templates, the library template were synthesized by Integrated DNA Technologies (IDT). The D-RNA aptamers containing both 2'-F-pyrimidines and native pyrimidines, 5'-FAM-labeled RNAs, 5'-biotin-labeled RNAs, all L-RNAs as aptamers and binding targets, were synthesized in house. The L-nucleoside phosphoramidintes were purchased from Chemgenes. The 2'-F-modified L-uridine and L-cytidine phosphoramidites were synthesized following our lab's reported protocols. Streptavidin Magnetic Beads (New England Biolabs) were used in capturing bound RNAs. T7 in vitro transcription was performed using the in-house expressed Y639F T7 polymerase mutant. Reverse transcription was performed using the ProtoScript™ II Re-verse Transcriptase from NEB. PCR amplification of the enriched library was performed using the Q5 High-Fidelity DNA Polymerase from NEB.

**1b. RNA oligonucleotides synthesis.** RNA oligonucleotides were synthesized by standard solid-phase phosphoramidite chemistry on a ABI RNA/DNA oligonucleotide synthesizer. Cleavage and elution of the full-length products from 1 μmol universal CPG-solid support columns, as well as removal of protecting groups on the nucleobases and phosphates, was carried out by heating the solid support for 15 min at 65 °C. The resultant homogeneous mixtures were first concentrated under reduced pressure for 2 hrs on a Genevac EZ-2 table top speedvac system (Genevac, Stone Ridge, NY), then lyophilized to dryness on a Labconco Benchtop 4.5 L freeze-drier (Labconco, Kansas City, MO) at <200 mTorr overnight to afford off-white solid residues. The residues were then resuspended in 115 μL of DMSO. 75 μL of TEA and 65 μL of TEA·3HF were added, and the solutions were heated for 2.5 hr at 65 °C to remove the TBDMS protecting group on the ribose 2'-hydroxyl group. The mixtures were homogeneous and pale to golden yellow in color. After cooling to room temperature (~30 mins), the RNA sample was desalted and detritylated by using the Glen-Pak RNA purification cartridge, following the protocol from Glen Research Inc. The purification of the desired products was performed by running denaturing polyacrylamide gels (urea-PAGE). The gel pieces containing the RNA fractions were collected, fragmented, and soaked in water for 12 hours. The RNA samples were then desalted by RNA precipitation using sodium acetate and 1-butanol. The precipitates were spun down (10000 rpm, 30 mins) and supernatants were removed by decanting. The resulting white solids were washed twice with absolute ethanol; the samples were then dried under high vacuum overnight before dissolving into water for appropriate concentrations. MALDI-TOF analysis of the SELEX target, 5'-biotin-labeled L-RNA attenuator hairpin, was shown in Figure S1.

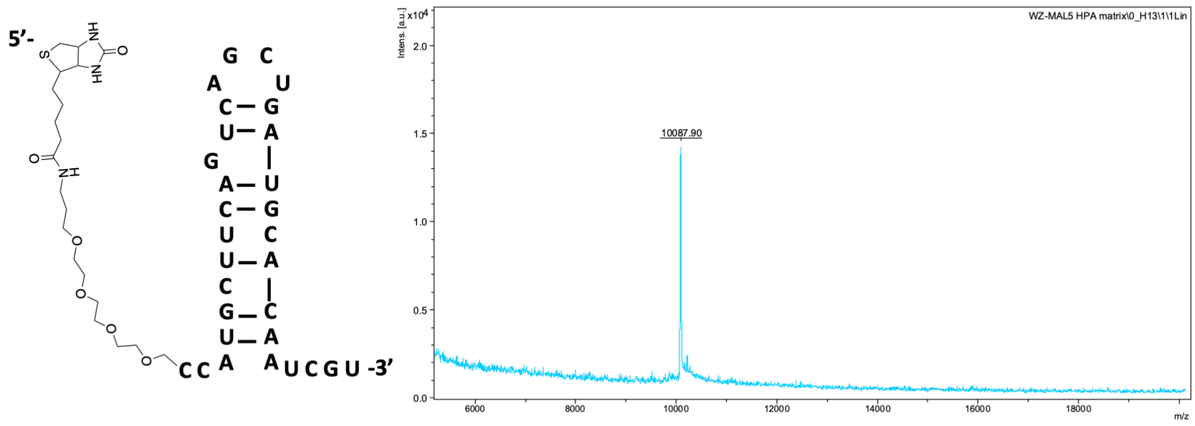


Figure S1. Sequence and MALDI-TOF result of the synthetic SELEX target, 5'-biotin-L-AH. Measured 10087.9 (calcd. 10061.4)

### 2. Protocols in SELEX and Enzymatic Reactions

**2a. In vitro SELEX.** Prior to each round, the RNA pool (1 nmol in the first round, in a mixture containing 50 mM NaCl, and 25 mM Tris pH 7.5) was heated at 75 °C for 1 min, then slowly cooled to 23 °C. An equal volume of another solution (20 mM $MgCl_2$, 50 mM NaCl, 25 mM Tris (pH 7.5), and 0.2% TWEEN-20) was added and mixed well. The combined mixture was incubated with Dynabeads that had been pre-blocked with yeast tRNA and washed (pre-block beads: 0.1 ug/µL Yeast tRNA, r.t. 2 hours, followed by washing 2x for equilibration into the binding buffer prior to use). The beads were discarded to remove bead-binding sequences from the RNA pool, then the 5'-Biotin labeled L-attenuator hairpin was added to the supernatant. The mixture was allowed to incubate at 23 °C for 60 min before adding Dynabeads. After shaking at room temperature for certain time, the beads were washed with a 500 uL washing solution containing 10 mM $MgCl_2$, 50 mM NaCl, 25 mM Tris (pH 7.5), and 0.1% TWEEN-20. Then the bound RNAs were eluted with two 200-μL of 25 mM NaOH. The eluted material was neutralized with 1 M Tris (pH 7.5) and purified using NEB RNA clean and concentration column. Selection pressure was increased over successive rounds by changing RNA concentrations, progressing lengthening the time of the washing steps, and changing the negative and positive selection times.

Table S1. Reaction conditions in each round of SELEX.

| ***Rounds*** | ***1*** | ***2*** | ***3*** | ***4*** | ***5*** | ***6*** | ***7*** | ***8*** | ***9*** |
|---|---|---|---|---|---|---|---|---|---|
| **RNA library conc. (uM)** | **5** | **2** | **1** | **0.5** | **0.5** | **0.2** | **0.1** | **0.1** | **0.1** |
| **Target conc. (uM)** | **2** | **1** | **0.5** | **0.5** | **0.2** | **0.1** | **0.1** | **0.05** | **0.05** |
| **Negative selection time (min)** | **5** | **10** | **20** | **30** | **30** | **45** | **45** | **60** | **60** |
| **Incubation time in positive selection (min)** | **60** | **60** | **60** | **45** | **45** | **45** | **30** | **30** | **20** |
| **Washing time in positive selection (min)** | **1** | **1** | **2** | **2** | **10** | **10** | **20** | **20** | **20** |
| **$MgCl_2$ conc. (mM)** | **10** | **10** | **10** | **10** | **10** | **10** | **10** | **10** | **10** |
| **NaCl conc. (mM)** | **50** | **50** | **50** | **50** | **50** | **50** | **50** | **50** | **50** |

**2b. T7 RNA Polymerase Mutant Expression and Purification** T7 RNA polymerase mutant (Y639F[1]) construct was a gift from Dr. Bin Zhu lab at Huazhong University of Science and Technology, China. Protein expression was carried out in BL21 (DE3) pLysS E. coli cells at 37°C in LB medium. Transformation was accomplished by heat shock method. A single colony was picked up to LB-ampicillin broth (20 mL). The culture was shaken (220 rpm) over-night at 37 °C. One liter LB-ampicillin-chloramphenicol broth was prepared and the overnight culture was added to inoculate it. When the OD600 of inoculated broth reached at 0.7 OD, protein expression was then induced by adding IPTG (final concentration 1 mM) and the culture was shaken (220 rpm) overnight at 20 °C. Cells were harvested by centrifugation and then lysed by soni-cation.

T7 RNA polymerase was purified on a Ni column and eluted in 20 mM Tris-HCl (pH 7.5), 300 mM NaCl, 5% glycerol, 1.4 mM β-mercaptoethanol, and 300 mM imid-azole. The buffer was exchanged to 20 mM Tris-HCl, 100 mM NaCl, 5% glycerol, 1.4 mM β-mercaptoethanol, and 0.5 mM EDTA for thrombin digestion. The activity of the protein was tested by the RNA transcription using 2'-fluoro-modified pyrimidines as substrates (Figure S2).

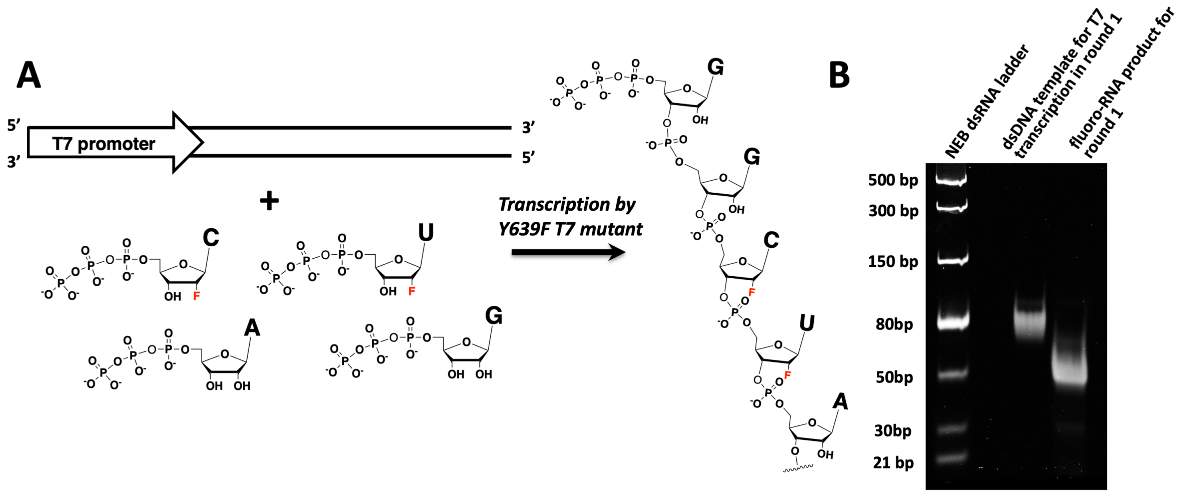


Figure S2. (A) Schematic representation of RNA transcription using Y639F T7 polymerase mutant. (B) PAGE gel analysis of transcription product in round 1.

**2c. RNA Transcription by Y639F T7 RNA Polymerase Mutant.** Double stranded template was used in a 100-μL transcription reaction containing T7 RNA polymerase (20 unit/μL), $MgCl_2$ (16 mM), DTT (10 mM), and 4 mM of each of ATP, GTP, 2'-F-UTP and 2'-F-CTP. Reaction solution was incubated at 37 °C for 14 h, before adding 2 U/μL DNase I. After an additional 30min at 37 °C, the mixture was purified by denaturing PAGE gel. The purified RNA pool was desalted by ethanol precipitation or cartridge. Before SELEX, the purity is monitored by analytical PAGE.

**2d. Reverse Transcription.** In each round, the eluted aptamers were reverse transcribed in a 40-μL reaction mixture containing 10 U/μL reverse transcriptase, 1 μM Reverse primer, and 1 mM of each of the four dNTPs. The RNA template/primer/dNTP complex in water was quickly annealed at 65°C then on ice to destruct aptamer's secondary structure. After 4 h at 42 °C of reverse transcription, 10 μL 1 M NaOH was added and incubated at 95°C for 10 min to denature the enzyme and degrade the unwanted RNA template. The mixture was neutralized by the addition of 30 μL of 1 M Tris-HCl (pH 7.6) before QIAGEN PCR Cleanup Kit.

**2e. PCR Amplification.** In each round, the cDNA was amplified by Q5 High-fidelity polymerase in a 100 μL PCR reaction, containing, 0.025 U/μL Q5 enzyme, 0.8 μM Forward primer, 0.8 μM Reverse primer, 0.2 mM of each of the four dNTPs, 1 mM $MgCl_2$. Hot-start PCR using, in a buffer containing Number of PCR cycles and the template quantity need to be optimized. Run 6% native PAGE to purify the PCR product if needed.

**2f. Microscale thermophoresis (MST) binding assays.** MST experiments were conducted in triplicate on a Monolith NT.115 system (NanoTemper Technologies). The RNA aptamer solutions with different concentrations were prepared in solutions containing 25 mM tris buffer (pH 7.5) and 50 mM NaCl. The RNA aptamer solutions were annealed by heated to 80°C for 2 min and then cooled to room temperature over 2 h, and then mixed with the synthetic 5'-FAM-labeled RNA attenuator hairpin, which was dissolved in a solution of 25 mM tris buffer (pH 7.5), 50 mM NaCl and 20 mM $MgCl_2$. 100 nM 5'-FAM-labeled RNA attenuator hairpin sample was used in the MST assays, and a 2-fold dilution series of aptamer was prepared, with the final concentrations of aptamers ranging from 100 μM to 3.1 nM. Samples were incubated on ice for at least 2 hours. Following incubation, the samples were added to premium coated capillaries (NanoTemper Technologies) and subsequently subjected to MST analysis at 25oC using blue light mode from the binding software. The results were analyzed by MST nano temper analysis (nta) analysis software using the Kd mode analysis, following the single-binding site model.

**2g. Confocal microscopy imaging of internalization of L-RNA aptamer.** Cells were grown on poly-d-lysine-coated glass cover slides in normal growth media and allowed to attach overnight. Cells were washed once with Opti-MEM medium (Thermo, Cat. #11058021) prior to treatment with Cy5 labelled aptamer mixed with Lipofectamine 3000 transfection agent (Thermo Fisher, Cat. # L3000001).

For confocal microscopy, cells were fixed by 4% paraformaldehyde, permeabilized using PBS-T (0.01% Triton X) and stained by Alexa Fluor® 488 phalloidin (Invitrogen) for cytoskeleton and DAPI for nucleus. Confocal microscopy was performed using a

Zeiss AxioObserverZ1 modified by 3i for confocal microscopy with the addition of a CSU-X1 M1 Spinning DiskConfocal and a Prime BSI CMOS Detector.

**2h. NF-κB Activation Assay in HEK293t Cells.** HEK293T cells were cultured in DMEM with 10% fetal bovine serum and 1% penicillin-streptomycin at 37°C with 5% $CO_2$. Cells were seeded at a density of $2\times10^5$ cells per well in 6-well plates and treated the next day with the RNA aptamers at 5 μM concentration for 12 hours. After treatment, cells were lysed using RIPA buffer supplemented with protease and phosphatase inhibitors. Proteins were extracted for Western blot analysis using the primary antibody against phospho-NF-κB p65 (Thermo Fisher, Catalog #MA5-15160), followed by a secondary antibody conjugated to horseradish peroxidase (HRP). β-actin served as a loading control. Band intensities were visualized and quantified by Bio-rad Chemidoc Imager by converting to relative light units (RLU).

Table S2. Enriched sequences from round 8, 9 and 10.

| *Name* | *Sequence* | *Kd (μM)* | *Percentage in pool* |
|---|---|---|---|
| 8.1 | GGU UAC CAG CCU UCA CUG CAU GUA CUG CUU GCU ACG GAA CGA ACG UGC GGU AGC GAU UCU GUU ACG ACC ACG GUC GGU CAC AC | | 3.2 % |
| 8.2 | GGU UAC CAG CCU UCA CUG CAU UGA AUA CUA GAA AGG GAU GUG GUA GUU GUC CUU GUG UCG GAC AGC ACC ACG GUC GGU CAC AC (same as 9.2) | 2.5 ± 0.4 | 3.1 % |
| 8.3 | GGU UAC CAG CCU UCA CUG CUU GAU CCG CGA UUG UGA CGU GGU UUC UUG CUG UGU UGC GCU CGU UGC ACC ACG GUC GGU CAC AC (same as 9.3) | 2.1 ± 0.5 | 3.0 % |
| 8.4 | GGU UAC CAG CCU UCA CUG CAU CAA CUG CAU GCU AUC UUA CUG GAG UCU GGU GCA GAA UGC CUU UGC ACC ACG GUC GGU CAC AC | | 2.8 % |
| 8.5 | GGU UAC CAG CCU UCA CUG CAU GAA CAG CUU GCU UGC GAA GGG AGG UGU GGU GGU GUU UGU GAU AGC ACC ACG GUC GGU CAC AC (same as 9.1) | 1.4 ± 0.2 | 1.6 % |
| 8.6 | GGU UAC CAG CCU UCA CUG CUU CCA AAG CAU ACU UGC GUA GCG CCG AGU CCU AGC ACU ACU CUA CCC ACC AGG GUC GGU CAC AC | | 1.2 % |
| 9.1 | GGU UAC CAG CCU UCA CUG CAU GAA CAG CUU GCU UGC GAA GGG AGG UGU GGU GGU GUU UGU GAU AGC ACC ACG GUC GGU CAC AC | 1.4 ± 0.2 | 10.8% |
| 9.2 | GGU UAC CAG CCU UCA CUG CAU UGA AUA CUA GAA AGG GAU GUG GUA GUU GUC CUU GUG UCG GAC AGC ACC ACG GUC GGU CAC AC | 2.5 ± 0.4 | 9.7% |
| 9.3 | GGU UAC CAG CCU UCA CUG CUU GAU CCG CGA UUG UGA CGU GGU UUC UUG CUG UGU UGC GCU CGU UGC ACC ACG GUC GGU CAC AC | 2.1 ± 0.5 | 8.5% |
| 9.4 | GGU UAC CAG CCU UCA CUG CUU CUG UAC UUC UAA UUC AUU AUU ACC UAC CUC GUG CUU UGU GCA CGC ACC ACG GUC GGU CAC AC | n.d. | 5.0% |
| 9.5 | GGU UAC CAG CCU UCA CUG CAU UCG AGA UAC AUG UGA UAG UAU CUC CUG AUG AUG GCU UUC GAC AGC ACC ACG GUC GGU CAC AC | n.d. | 4.2% |
| 9.6 | GGU UAC CAG CCU UCA CUG CAU UGU GGG GCC UUU AGU GAU UAA GGC CAU GCA CUG UUG GCU UGU GGC ACC ACG GUC GGU CAC AC | n.d. | 3.3% |
| 10.1 | GGU UAC CAG CCU UCA CUG CAU UGA AUA CUA GAA AGG GAU GUG GUA GUU GUC CUU GUG UCG GAC AGC ACC ACG GUC GGU CAC AC (same as 9.2) | 2.5 ± 0.4 | 11.8 % |
| 10.2 | GGU UAC CAG CCU UCA CUG CAU GAA CAG CUU GCU UGC GAA GGG AGG UGU GGU GGU GUU UGU GAU AGC ACC ACG GUC GGU CAC AC (same as 9.1) | 1.4 ± 0.2 | 10.6 % |
| 10.3 | GGU UAC CAG CCU UCA CUG CAU UGU GGG GCC UUU AGU GAU UAA GGC CAU GCA CUG UUG GCU UGU GGC ACC ACG GUC GGU CAC AC (same as 9.6) | | 8.3 % |
| 10.4 | GGU UAC CAG CCU UCA CUG CUU GAU CCG CGA UUG UGA CGU GGU UUC UUG CUG UGU UGC GCU CGU UGC ACC ACG GUC GGU CAC AC (same as 9.3) | 2.1 ± 0.5 | 7.6 % |
| 10.5 | GGU UAC CAG CCU UCA CUG CAU GUA CAG CAU GCU ACG GCA UGC AGU UGC CCU GCA CAU GCA GCA AGC GCC AUG GUC GGU CAC AC | | 3.1 % |
| 10.6 | GGU UAC CAG CCU UCA CUG CAU GUA CUG CUU GUU CAG AUU GCU AGG ACG UUG AAC CAA CCC ACU AGC AGC ACC GUC GGU CAC AC | | 2.0 % |

n.d.= detectable

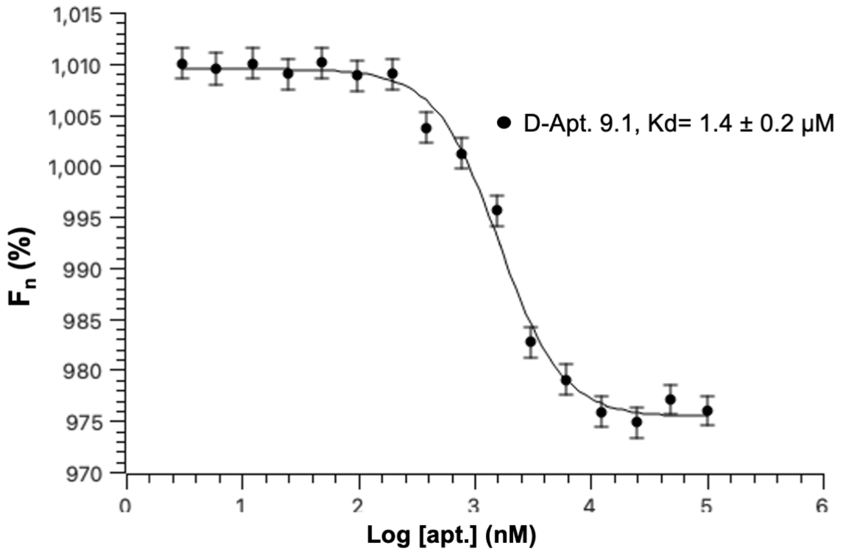


Figure S3. Binding curve in MST assays generated from the difference in initial fluorescence intensity compared to the intensity in the presence of heat (D-Apt. 9.1 against FAM-L-AH).

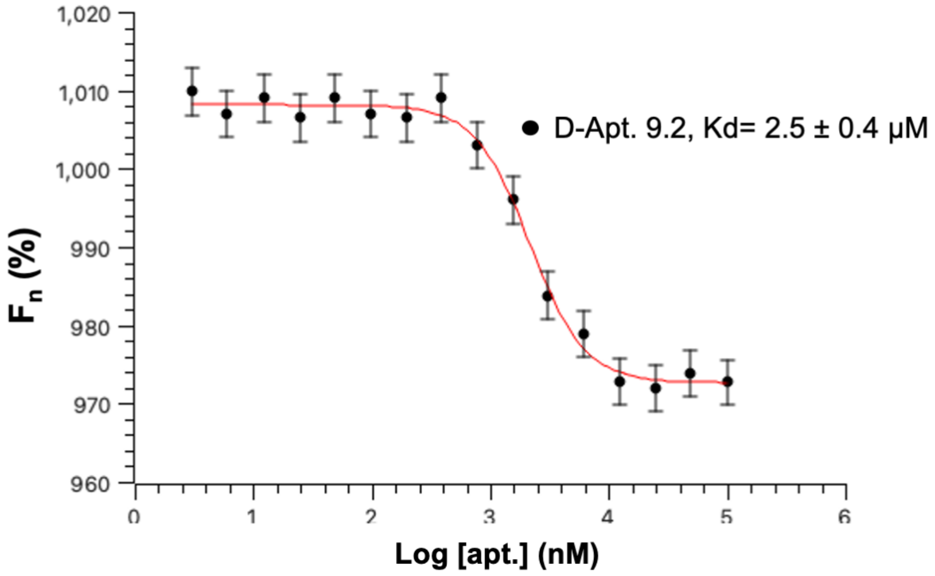


Figure S4. Binding curve in MST assays generated from the difference in initial fluorescence intensity compared to the intensity in the presence of heat (D-Apt. 9.2 against FAM-L-AH).

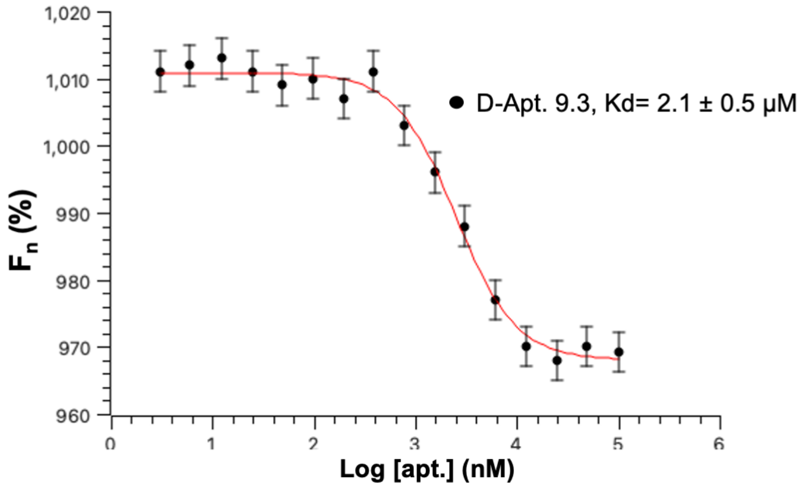


Figure S5. Binding curve in MST assays generated from the difference in initial fluorescence intensity compared to the intensity in the presence of heat (D-Apt. 9.3 against FAM-L-AH).

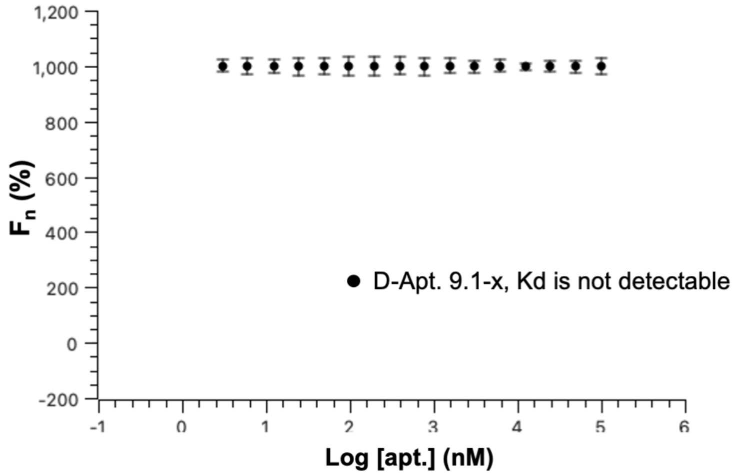


Figure S6. Binding curve in MST assays generated from the difference in initial fluorescence intensity compared to the intensity in the presence of heat (D-Apt. 9.1-x against FAM-L-AH).

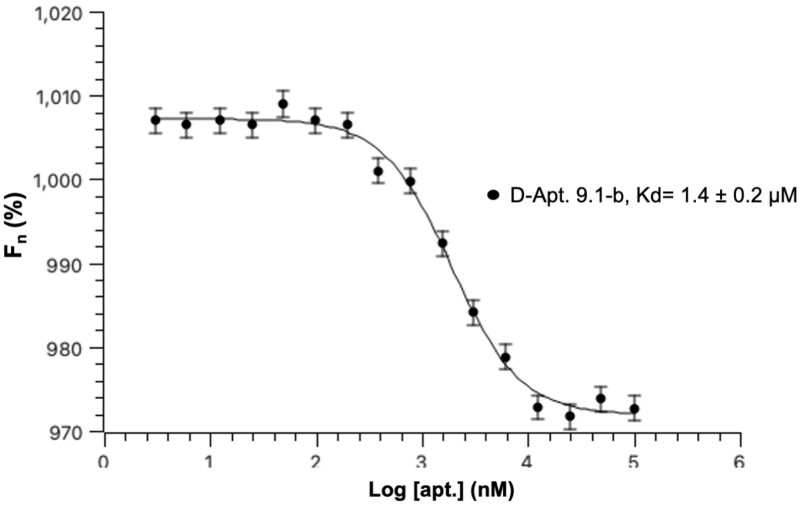


Figure S7. Binding curve in MST assays generated from the difference in initial fluorescence intensity compared to the intensity in the presence of heat (D-Apt. 9.1-b against FAM-L-AH).

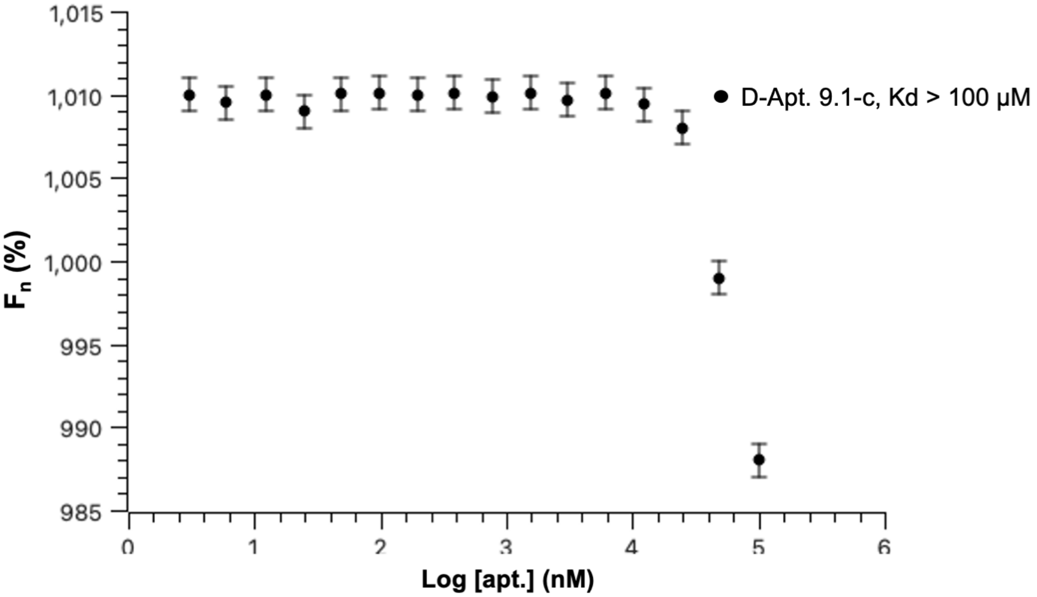


Figure S8. Binding curve in MST assays generated from the difference in initial fluorescence intensity compared to the intensity in the presence of heat (D-Apt. 9.1-c against FAM-L-AH).

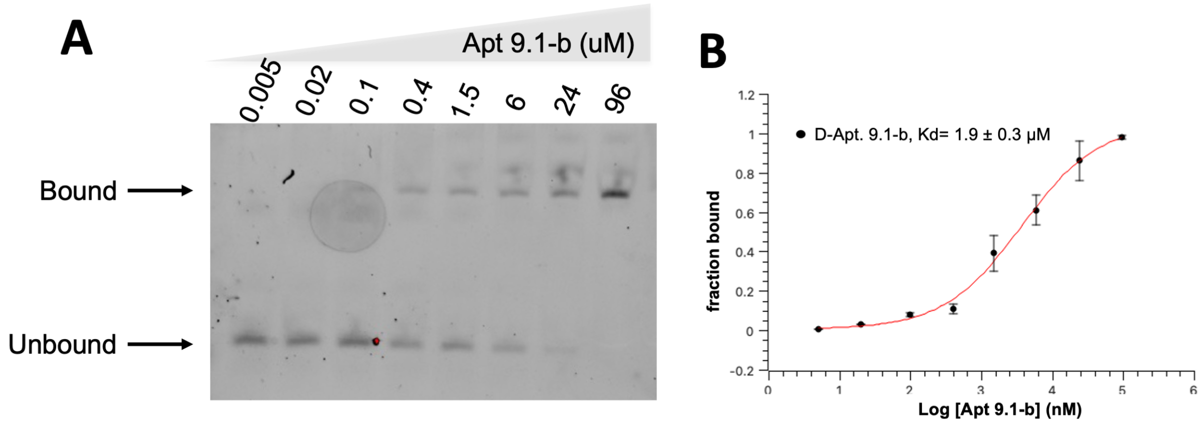


Figure S9. EMSA analysis of Apt 9.1-b binding to L-AH target. (A) Gel shift experiment using FAM-labeled L-AH and increasing concentration of Apt 9.1-b. (B) Binding curves of D-Apt. 9.1-b against FAM-L-AH. Kd is found to be 1.9 ± 0.3 μM.

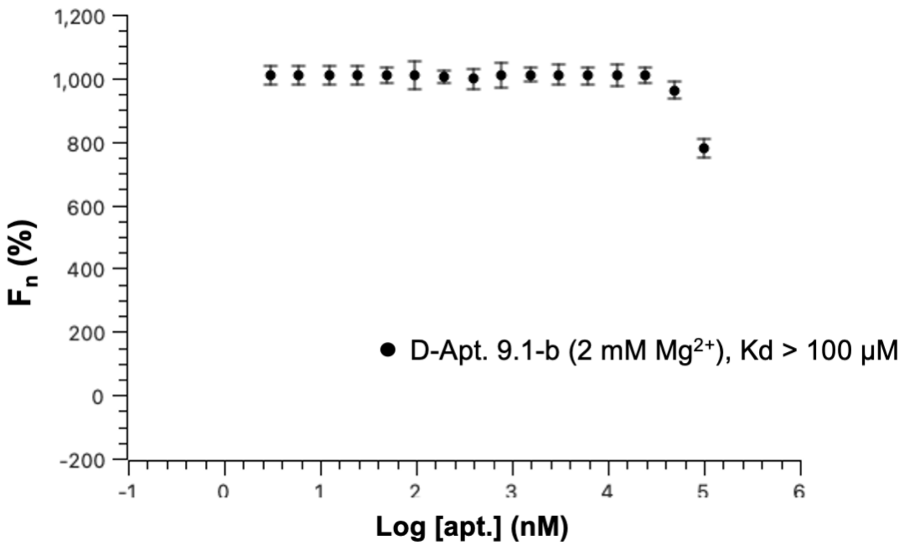


Figure S10. Binding curve in MST assays generated from the difference in initial fluorescence intensity compared to the intensity in the presence of heat (D-Apt. 9.1-b against FAM-L-AH at 2 mM $Mg^{2+}$).

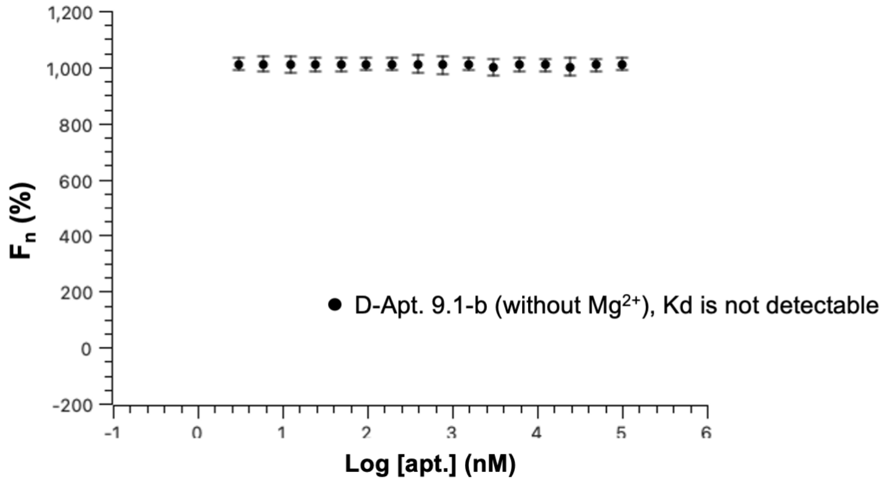


Figure S11. Binding curve in MST assays generated from the difference in initial fluorescence intensity compared to the intensity in the presence of heat (D-Apt. 9.1-b against FAM-L-AH without $Mg^{2+}$).

### 3. References.


(1) Padilla, R.; Sousa, R. A Y639F/H784A T7 RNA polymerase double mutant displays superior properties for synthesizing RNAs with non-canonical NTPs. *Nucleic Acids Res.* **2002**, *30* (24), e138.
